# Thermal Tolerance Plasticity and Dynamics of Thermal Tolerance in *Eublepharis macularius*: Implications for Future Climate-Driven Heat Stress

**DOI:** 10.1101/2024.01.13.575519

**Authors:** Emma White, Solyip Kim, Garrett Wegh, Ylenia Chiari

## Abstract

The intensity and duration of heat waves, as well as average global temperatures, are expected to increase due to climate change. Heat waves can cause physiological stress and reduce fitness in animals. Species can reduce overheating risk through phenotypic plasticity, which allows them to raise their thermal tolerance limits over time. This mechanism could be important for ectotherms whose body temperatures are directly influenced by available environmental temperatures. Geckos are a large, diverse group of ectotherms that vary in their thermal habitats and times of daily activity, which could affect how they physiologically adjust to heat waves. Data on thermal physiology are scarce for reptiles, with only one study in geckos. Understanding thermal tolerance and plasticity, and their relationship, is essential for understanding how some species are able to adjust or adapt to changing temperatures. In this study, we estimated thermal tolerance and plasticity, and their interaction, in the crepuscular gecko, *Eublepharis macularius*, a species that is emerging as a model for reptile biology. After estimating basal thermal tolerance for 28 geckos, thermal tolerance was measured for each individual a second time at several timepoints (3, 6, or 24 h) to determine thermal tolerance plasticity. We found that thermal tolerance plasticity (1) does not depend on the basal thermal tolerance of the organism, (2) was highest after 6 hours from initial heat shock, and (3) was negatively influenced by individual body mass. Our findings contribute to the increasing body of work focused on understanding the influence of biological and environmental factors on thermal tolerance plasticity in organisms and provide phenotypic data to further investigate the molecular basis of thermal tolerance plasticity in organisms.

## 1. Introduction

Anthropogenic climate change has led to the rise of extreme heat events caused by higher average temperatures across the year (Stillman et al., 2019). Extreme heat events such as heat waves have become more prevalent within the summer months and are expected to increase in frequency, duration, and severity in the coming century (Stillman et al., 2019; Angélil et al., 2017; Guo et al., 2018; Pachauri et al., 2014). Heat waves can induce physiological stress on organisms and can reduce fitness (Stillman et al., 2019; Coffel et al., 2017; Froelicher et al., 2019). This is especially worrisome for species living in warm, aseasonal environments, since these species typically have narrow thermal tolerance ranges compared to species living in more variable, heterogeneous environments (Stillman et al., 2019, Williams et al., 2016; Tewksbury et al., 2008; Hoffmann et al., 2012; Ruibal, 1961; Janzen, 1967). Organisms can respond in many ways to high and potentially lethal temperatures; for example, behaviorally by seeking cooler microclimates within their local environment to prevent overheating (Stillman et al., 2019). At the same time, organisms may also plastically adjust the upper thermal tolerance of their internal body temperature when exposed to thermal extremes. This process of physiological plasticity is mediated by relatively rapid gene expression and cellular response changes (Kregel, 2002; Hu et al., 2014; Hamdoun et al., 2002; Horowitz, 2001; Buckley & Hofmann, 2002; Stillman et al., 2019, Angilletta, 2009), so that organisms plastically adjusting the upper limit of thermal tolerance can revert back to the original state after the heat stress has occurred, for example in response to short thermal perturbations (Sørensen et al., 2019).

Rapid reversible thermal tolerance plasticity could happen through heat hardening. Heat hardening (Bowler, 2005) occurs when an organism is exposed to its thermal extreme (either hot or cold) over a short period of time (minutes to hours), triggering a stress response in the organism with a consequent production of an abundance of heat-shock proteins (HSPs) or cold-shock proteins (CSPs) (Horowitz, 2001; Sørensen et al., 2003; Gangloff & Telemeco, 2018). HSPs act as molecular chaperones, protecting other cellular proteins from denaturing at higher temperatures and allowing the organism to increase its thermal tolerance over time (Kregel, 2002; Daugaard & Jäättelä, 2007; Sørensen et al. 2003; Gangloff & Telemeco, 2018). HSP expression has been shown to correlate with thermal tolerance across ectotherm species, where species adapted to warm environments (i.e., desert) have higher HSP concentrations and maintain the production of HSPs at higher temperatures compared to species adapted to cooler environments (i.e., forest) (Ulmasov et al., 1992; Zatespina et al., 2000; Madeira et al., 2012; Madeira et al., 2014; Qin et al., 2003; Newman et al., 2005; Dehghani et al., 2011). Heat hardening capacity varies across ectotherm species and populations (Phillips et al., 2016; Deery et al., 2021; Gilbert & Miles, 2019; Lapwong et al., 2021; Mottola et al., 2022; Morgan et al., 2018; van Heerwaarden et al., 2014). The heterogeneity of the thermal conditions during the developmental stages of an organism and of the habitat in which the species occurs (i.e., seasonal thermal variation and differences across the distribution range of the species), for example, are known to often influence heat hardening: the more variable the thermal environment is, the more plastic the organism will be in its response to thermal variation (Manenti et al., 2014; Phillips et al., 2016; Deery et al., 2021; Gilbert & Miles, 2019; Sasaki & Dam, 2019; but see Gunderson & Stillman, 2015; van Heerwaarden et al., 2014). Heat hardening is also strongly influenced by the biology of the species (Gunderson et al., 2017; Deery et al., 2021). For example, in ectotherms, the internal body temperatures, and thus the physiology and behavior of these organisms, are directly affected by temperatures available within their respective environments (Huey, 1991; Dayananda et al., 2016). As such, ectotherms can be particularly vulnerable to thermal extremes and thus thermal tolerance plasticity through heat hardening has the potential to reduce overheating risks.

In addition to the influence of thermal environmental heterogeneity, the “trade-off” hypothesis (TOH) has been proposed as another mechanism to explain variation in heat hardening across and within species (van Heerwaarden & Kellermann, 2020). This hypothesis suggests that species already adapted to high temperatures, thus having high thermal tolerance, have limited potential to improve their thermal tolerance via phenotypic plasticity. The consequence of this hypothesis is that species cannot evolve both high thermal tolerance and high plasticity in response to extreme temperatures (van Heerwaarden & Kellermann, 2020). The TOH has recently been debated to be supported across and within different lizard species because not all species show a negative correlation between heat hardening capacity and basal upper thermal tolerance (Gunderson, 2023), which suggests that other factors may be influencing variation in thermal tolerance plasticity. Data on heat hardening are available for an extremely limited number of lizard species (only six species out of the 7,641 species described, Uetz et al., 2023) (Phillips et al., 2016; Deery et al., 2021; Gilbert & Miles, 2019; Lapwong et al., 2021).

Among lizards, geckos - a clade including 1,935 species (Uetz et al., 2020) - are tremendously understudied with data so far available for only one species (Lapwong et al., 2021). However, geckos have a wide range of variation for traits that can influence their thermophysiology and potentially their thermal tolerance, including time of activity, habitat, distribution, locomotion, morphology, and body size (Meiri, 2019; Gamble et al., 2015; Grismer et al., 2015; Heinicke et al., 2017; Oliver et al., 2019). As nighttime air temperatures have been increasing at a faster rate than daytime air temperatures due to climate change (Karl et al., 1991; Easterling et al., 1997; Vose et al., 2005; Alexander et al., 2006), understanding how species active during the crepuscular hours or at night can respond to increasing temperatures – something largely unknown across squamate reptiles – will allow predictions on the influence of climate change on the fitness of species active in dim light (Rutschmann et al., 2023).

In this study, we established the critical thermal maximum (basal upper thermal tolerance), inferred the thermal tolerance plasticity and assessed their relationship in the gecko, *Eublepharis macularius. E. macularius* is becoming a model species for reptile studies in various fields of research (e.g., Hastings et al., 2023, Pinto et al., 2023, Katlein et al., 2022, Glimm et al., 2021, Kiskowski et al., 2021) and has well-documented molecular and genomic data compared to most other lizard species. *E. macularius* is typically active during dusk (Khan 2009; Gamble et al., 2015) and native to parts of northeast Iran and northwest India, where it inhabits deserts, shrublands, grasslands and rocky cliffs (Papenfuss et al., 2021). In the coming century, the native habitat of *E. macularius* is expected to experience higher frequency and duration of heat waves (Rohini et al., 2019; Khan et al., 2020), which could increase this species’ vulnerability to heat stress (Stillman et al., 2019).

In ectotherms, thermal tolerance plasticity through heat hardening has been measured as the change in the critical thermal maximum (CT_max_) over time. CT_max_ corresponds to the upper thermal tolerance of an organism, which is the internal body temperature at which the organism loses physiological function and cannot tolerate any higher temperatures (Angilletta, 2009). Measurements of CT_max_ can be influenced by heating rates (i.e., the rate at which individuals approach their upper thermal tolerance) (Ribeiro et al., 2012; Terblanche et al., 2007), which have been reported in previously published heat hardening studies; however, validation of consistent heating rates using statistical analyses is generally missing from methodologies in these papers. As such, in addition to measuring heat hardening capacity and thermal tolerance within *E. macularius*, in this work we also investigate the influence of individual body mass, body size, and sex on heating rates across CT_max_ experiments. We hypothesized that: (1) in support of the TOH, individuals with lower basal thermal tolerance have greater heat hardening capacity compared to individuals with higher basal thermal tolerance; (2) overtime, change in thermal tolerance will be highest at 6 hours after initial heat shock, following what has been shown in other lizard species (Phillips et al., 2016, Lapwong et al., 2021; Gilbert & Miles, 2019); and (3) individuals with larger body size will have greater heat hardening capacity compared to smaller individuals since acclimation capacity has been shown to increase with body size across ectotherms (Rohr et al., 2018).

Ultimately, the costs and benefits of heat hardening for organisms will be dependent upon the thermal extremes (either hot or cold) that an organism is exposed to within its native environment (Stillman et al., 2019; Noer et al., 2022). Although, heat hardening alone may not allow organisms to fully compensate for the effects of climate warming in the future (Gunderson et al., 2017), it has been estimated to reduce overheating risk in species to some degree (Gunderson et al., 2017; Deery et al., 2021). As such, understanding the breadth of thermal tolerance plasticity for an organism gives context for how the organism may respond to future temperature extremes in their native environment and whether the organism can physiologically adjust quickly enough to the thermophysiological stress associated with these higher environmental temperatures.

## 2. Materials and Methods

All capture, handling, and experimental protocols were approved by George Mason University IACUC committee (Permit number #1901361). Experiments were carried out to minimize stress and disturbance to the animals and in accordance with relevant guidelines and regulations.

### 2.1 Study species and captivity conditions

A total of 28 adult individuals (14 females and 14 males) were used for this study. Only adult individuals of *E. macularius* were tested, as using individuals of different life-stages may affect measurements of critical thermal maximum (CT_max_) (Telemeco & Gangloff, 2021; Camacho & Rusch, 2017). All geckos were part of a captive colony housed at George Mason University for several years. In their captive conditions, geckos were housed in plastic cages (Sterilite 28 quart/27 liter clear storage box) with two small plastic bowls of which one is for food and one for water, humid and dry hide boxes, and heating pads at one end of the cage for thermoregulation. In their captive environment, geckos were fed every other day with a mixed diet of mealworms, crickets, and waxworms dusted with calcium powder and had access to water *ad libitum.* Gecko cages were housed in a temperature-controlled room operating on a 12h:12h light-dark cycle with daily room temperatures ranging from 24°C to 26°C and relative humidity ranging from 20-50%. All individuals were obtained from different captive breeders and held in these laboratory conditions for at least two months before the start of CT_max_ experiments.

### 2.2 Experimental set up

Prior to conducting the experiments, geckos were fasted for at least three days to prevent any influence of feeding on CT_max_ measurements. Before the start of each experiment (both basal and final CT_max_ for each individual), the mass (g) of each individual was measured using a digital scale. In addition, each individual’s snout-vent-length (SVL) was measured for all geckos. Final CT_max_ corresponds to the CT_max_ obtained to estimate heat hardening and measured at 3h, 6h, or 24h.

CT_max_ experiments were performed at George Mason University in an empty temperature-controlled room set to 28°C, which enabled individuals to acclimate to an internal body temperature of 28°C prior to starting CT_max_ experiments. For CT_max_ experiments, individuals were placed in a plastic holding container (Hefty 6.5 quart) placed on a lab bench in the temperature-controlled room with the 150W heat emitter domed lamp (ReptiZoo, 120V, 60Hz) placed 23 cm above the base of the container using a lamp stand. All sides of the plastic holding container were covered with Teflon, which was secured to the bottom of the plastic holding container using black electrical tape to ensure that the plastic holding container did not overheat during CT_max_ experiments. An iButton data-logger (DS1921G Thermochron, Maxim Integrated Products, precision = +/− 0.5°C) was placed both inside and outside the plastic holding container during CT_max_ experiments to monitor ambient temperatures (Supp. Materials Figure S1 for images of the experimental setup).

Geckos were tested one at the time. Unless a gecko was tested, it was in its housing room and cage. For all measurements of CT_max_, it is important that all individuals start at the same internal body temperature, as starting temperatures have been shown to influence CT_max_ values (Terblanche, et al., 2007; Camacho and Rusch, 2019). Prior to the start of CT_max_ experiments, a T-type 36-gauge wire thermocouple (Evolution Sensors and Controls, LLC) was inserted approximately 3 mm into the cloaca of the gecko to measure internal body temperature using a digital thermometer (Evolution Sensors and Controls, LLC; resolution = 0.1t < 1000° 1.0t ≥ 1000°; accuracy = ±0.1% + 0.6°C). If an individual’s cloacal temperature was approximately 28°C (± 0.9°C), CT_max_ experiments would be started. If an individual’s cloacal temperature was lower or higher than 28°C, the individual would either be placed in a water bath until the individual’s body temperature was lowered to 28°C, or in the plastic holding container to allow its internal body temperature to acclimate to 28°C as the room temperature. The small plastic container filled with water was used to allow the organism to recover after measurements of CT_max_.

### 2.3 Data collection

Basal CT_max_ - corresponding to the thermal tolerance of the individual at 0 hours - was measured for each of the 28 adult geckos as the internal body temperature at which each individual lost righting response for 10 sec. after being flipped over the individual’s dorsal side. This approach has been commonly used as a method of measuring CT_max_ in lizards (Deery et al., 2021, Phillips et al., 2016; Lapwong et al., 2021), as it is related to the internal body temperature at which the organism loses proper physiological function and cannot tolerate any higher temperatures.

To measure heat hardening (thermal tolerance plasticity) as the change in CT_max_ over time, the 28 individuals were randomly assigned a time interval to measure CT_max_ again after the initial measurement of basal CT_max_. Between measurements of basal and final CT_max_, individuals were taken from the testing room and placed in their respective housing enclosures to recover, since allowing the individual to remain at its basal CT_max_ would be lethal to the organism. Time intervals for measuring CT_max_ a second time (final CT_max_) ranged from 3 hours, 6 hours, or 24 hours after initial basal CT_max_ measurements. All individuals were initially tested at 0h for their basal CT_max_ and then at either 3h, 6h, or 24h after 0h (final CT_max_), depending on their assigned time-interval treatment group. These time intervals were chosen based on previously published studies using similar time intervals across 24 hours to measure CT_max_. These studies showed that heat hardening occurs as a short-term plastic response over 24 hours with the highest change in CT_max_ for most species occurring at 6 hours after initial heat shock (Deery et al., 2021; Lapwong et al., 2021; Phillips et al., 2016; Gilbert and Miles, 2019). The 3h time-interval group included 4 male and 5 female individuals, the 6h time-interval group included 5 male and 4 female individuals, and the 24h time-interval group included an equal ratio of 5 male and 5 female individuals.

Although *E. macularius* is a crepuscular gecko species, individuals would be expected to experience higher environmental temperatures during the day in this species’ native environment (Supp. Materials Figure S2 A and B). Thus, all CT_max_ experiments were conducted during daytime hours. Basal CT_max_ experiments were conducted from approximately 10:00 am to 12:00 pm, and final CT_max_ experiments were conducted 3 hours, 6 hours, and 24 hours after the start of basal CT_max_ experiments.

To obtain data on basal and final CT_max_, the internal body temperature of each individual for all CT_max_ experiments was raised at approximately 1°C/min (mean basal heating rate = 0.976 °C/min; mean final heating rate = 0.911°C/min) using a 150W heat emitter domed lamp placed above the experimental plastic container (Supp. Materials Figure S1) until each individual lost righting response for 10 seconds. Different heating rates can affect CT_max_ values, which are mainly affected by differences in individual body size (Ribeiro et al., 2012; Terblanche et al., 2007; Rezende et al., 2011). Because of this, a heating rate of 1°C/min – similar to what used in previous studies (Phillips et al., 2016; Gilbert & Miles, 2019; Lapwong et al., 2021) – was targeted for all individuals. To measure CT_max_ for each individual (basal and final CT_max_), individuals were flipped on their dorsal side inside the plastic holding container to check for righting response and cloacal temperature was measured immediately by quickly (within 10 seconds max) lifting the gecko from the plastic holding container and inserting 3 mm into the cloaca a T-type 36-gauge wire thermocouple attached to a digital thermometer (Evolution Sensors and Controls, LLC). Previous heat hardening experiments conducted on different lizard species left the thermocouple inside the cloaca during the entire CT_max_ experiment by securing it with a piece of medical tape (Deery et al., 2021). However, prior to conducting our trial experiments with individuals not used in this study, leopard geckos were able to remove the medical tape and the attached thermocouple from their cloaca and we could therefore not use this approach in our study. After measuring CT_max_ (both basal and final), the gecko was placed in a small container filled with water to allow the organism to recover.

### 2.4 Native microclimate temperatures for *E. macularius*

Seasonal variation of native environmental temperatures has been shown to influence heat hardening capacity in lizard species (Phillips et al., 2016). Therefore, in addition to measuring thermal tolerance and heat hardening capacity of *E. macularius*, both daily and seasonal temperature ranges of native microclimates were estimated for this species. To obtain an overall estimation of the native microclimatic environmental temperatures for *E. macularius*, occurrences for this species were downloaded from the Global Biodiversity Information Facility (GBIF, September 2023) and imported into Rstudio (v. 4.3.0, R Core Team 2023) using the “occ_download_get” function from the *rgbif* package (Chamberlain et al., 2017). Species occurrences for *E. macularius* were filtered for species’ scientific name mismatches as well as NA values for latitude, longitude, species’ scientific names, and country codes. Species occurrences were also cleaned and cross-checked for coordinate validity using the “clean_coordinates’’ function from the package, *Coordinate Cleaner* (Zizka et al., 2019). Species occurrences which resulted in at least one flagged test labeled as, “FALSE,” were removed from the dataset. From a total of 23 unique occurrences for *E. macularius*, microclimate temperatures were estimated using the “micro_global” function from the package *NiceMapR* (Kearney & Porter, 2017). Microclimate temperatures for each coordinate were estimated every 60 min across 365 days for 10 years using the NicheMapR microclimate model. Local height for the model was set to 1.5 cm, which is the measured midpoint height of captive *E. macularius*. In addition, the maximum shade parameter was set to 0% and 100% to estimate microclimate temperatures in full sun and full shade, respectively. Native microclimate temperatures estimated for *E. macularius* are shown in the Supplementary Materials (Supp. Materials Figure S2).

### 2.5 Statistical analyses

All statistical analyses were performed using the software Rstudio (v. 4.3.0, R Core Team 2023). All the tested 28 individuals were used for all analyses. A paired, two-sample t-test was first used to determine if the mass measured before basal CT_max_ and final CT_max_ experiments for all individuals significantly differed using the “t.test” function from the *stats* package (R Core Team 2023). We found a significant difference between mass measured before basal CT_max_ experiments and before final CT_max_ experiments across all individuals (t = 5.3729, df = 26, p-value = 1.259 x 10^-5^, mean difference = 0.911, 95% CI = (0.562, 1.259); as such, analyses were repeated taking into account the mass measured before basal CT_max_ or the mass taken before final CT_max_ experiments and tested independently. One-way ANOVAs were performed using the “aov” function from the *stats* package to determine if SVL or mass were influenced by the sex (male or females) of the individuals using separate models for each variable. One-way ANOVAs were also used to determine if SVL or mass significantly differed across time-interval treatment groups.

Individual heating rates during CT_max_ experiments were calculated as the slope corresponding to the change in internal body temperature over time measured every minute at time 0 (basal heating rate) and at the time of treatment (3, 6, or 24 hours). As such, each individual had a single basal and a single final heating rate value. The basal heating rate for each individual was calculated from the slope obtained by plotting the internal body temperatures obtained every minute during measurements of CT_max_ experiments at 0 hours on the time at which each measurement was obtained. Similarly, the final heating rate for each individual was calculated from internal body temperatures measured during CT_max_ experiments for either the 3h, 6h, or 24h time-interval treatment group. Paired, two-sample t-tests were performed to compare basal heating rates and final heating rates for all individuals within (e.g., for all the geckos tested after 3 hours) and across time-interval treatment groups (e.g., geckos tested at 3 hours, 6 hours, and 24 hours grouped together independently of the treatment time). The assumptions for each paired, two-sample t-test were checked using Shapiro-Wilks and Levene’s tests to examine the normality and homogeneity of variance for the differences between basal and final heating rates (Supp. Materials for details). In addition, one-way ANOVAs were performed to determine if basal heating rates for all individuals were influenced by time-interval treatment group, sex, mass measured before basal CT_max_ experiments, and SVL by testing each variable separately. Two-way ANOVAs were performed to test for the influence of sex, mass measured before basal CT_max_ and final CT_max_ experiments, and SVL separately on final heating rates with time-interval treatment group as a covariate for each model. To measure the potential influence of each individual on the fitted response values for each linear model as described above for basal and final heating rates, a Cook’s Distance test (Cook, 1977) was performed using the “cooks.distance” function on each model separately with a threshold of 1 (individuals with a Cook’s Distance > 1 are influential)

According to the TOH, individuals with low basal thermal tolerance are expected to have greater heat hardening and vice versa (van Heerwaarden & Kellermann, 2020). To test for a trade-off between basal thermal tolerance (basal CT_max_ for each individual) and heat hardening capacity (difference between basal CT_max_ and final CT_max_ for each individual), a Pearson’s product-moment correlation analysis was performed on the 3h, 6h, and 24h time-interval treatment groups using the “cor.test” function from the *stats* package. If the relationship between basal CT_max_ and change in CT_max_ (i.e., final CT_max_ at 3h, 6h, or 24h - basal CT_max_ at 0h) values (“unadjusted”) for each time-interval treatment group showed a significant (p-value < 0.05) negative relationship, then change in CT_max_ values were adjusted to remove the effect of regression to mean following the correction method by Kelly and Price (Kelly and Price, 2005). Change in CT_max_ values were only adjusted for the 3h time-interval treatment group and not for the 6h or 24h time-interval treatment group, since basal CT_max_ and change in CT_max_ showed a significant negative relationship (t-value = −4.8348, df = 7, p-value = 0.001889, r = −0.8772416, 95% CI = (−0.973, −0.510)) for the 3h time-interval treatment group. Following adjustment of change in CT_max_ values, the Pearson’s product-moment correlation analysis was then performed on the basal CT_max_ and adjusted change in CT_max_ values for the 3h time-interval treatment group.

The mean and standard deviation for CT_max_ values measured at either 0, 3, 6, or 24 hours were calculated using the “mean” (base R) and “sd” (*stats* package) functions, respectively. Standard error of measurement (SEM) was calculated for basal CT_max_ values measured at 0 hours or final CT_max_ values measured at either 3, 6, or 24 hours by dividing the standard deviation for each set of CT_max_ values by the square root of the sample size for each set of CT_max_ values. Generalized linear models (GLMs) were used to determine the effects of treatment groups on basal CT_max_ (basal CT_max_ at 0h) using sex, SVL, and mass as covariates in the models. GLMs for final CT_max_ (final CT_max_ at 3h, 6h, and 24h) and change in CT_max_ (final CT_max_ at either 3h, 6h, or 24h - basal CT_max_ at 0h) were also run using sex, SVL, mass, and the interaction between time-interval treatment group and basal CT_max_ as covariates in each of these models. The same GLMs for final CT_max_ and change in CT_max_ were also performed with the mass measured for each individual before final CT_max_ experiments instead of using the mass of each individual measured before basal CT_max_ experiments. Each GLM for basal CTmax, final CT_max_, and change in CT_max_ was then performed a second time with only significant variables included within each model. GLMs were performed using the “glm” function from the *stats* package. Change in CT_max_ included adjusted values for the 3h time-interval treatment group (see section above) and unadjusted values for the 6h and 24h time-interval treatment groups. The assumptions for each GLM were checked using Shapiro-Wilks and Levene’s tests to examine the normality and homogeneity of variance for basal CT_max_, final CT_max_, and change in CT_max_ values as well as the residual values for each GLM model (Supp. Materials for details). Linear regressions were used to determine the influence of basal CT_max_ on final CT_max_ separately within each time-interval treatment group (3h, 6h, and 24h) using the “lm” function from the *stats* package. To measure the influence of each individual on the fitted response values for all of the GLM models for basal CT_max_, final CT_max_, and change in CT_max_, a Cook’s Distance test was performed using the “cooks.distance” function on each model separately with a threshold of 1 (individuals with a Cook’s Distance > 1 are influential).

## 3. Results

### 3.1 Time-interval treatment group and sex differences

We found no significant differences in body mass measured before basal CT_max_ experiments (df = 1, sum of squares = 128.1, mean square = 128.14, F-value = 2.991, p-value = 0.0956) or final CT_max_ experiments (df = 1, sum of squares = 135.8, mean square = 135.83, F-value = 3.029, p-value = 0.0941) between male and female individuals; however, SVL (df = 1, sum of squares = 1.800, mean square = 1.8004, F-value = 5.426, p-value = 0.0279) significantly differed between sex with males being larger than females (male mean SVL = 12.06 cm; female mean SVL = 11.55 cm). Neither SVL (df = 2, sum of squares = 0.20, mean square = 0.0998, F-value = 0.244, p-value = 0.785), mass measure before basal CT_max_ experiments (df = 2, sum of squares = 35.2, mean of squares = 17.58, F-value = 0.364, p-value = 0.698) or mass measured before final CT_max_ experiments (df = 2, sum of squares = 47.4, mean square = 23.71, F-value = 0.47, p-value = 0.63) significantly differed across time-interval treatment groups.

### 3.2 Heating rates across and within time-interval treatment groups

Paired, two-sample t-tests found no significant differences between basal and final heating rates for individuals across all time-interval treatment groups (t = 1.7754, df = 27, p-value = 0.08711, mean difference = 0.0646, 95% CI = (−0.010, 0.139)) (Supp. Materials Figure S3), within the 3h (t = 0.36466, df = 8, p-value = 0.7248, mean difference = 0.0281, 95% CI = (−0.149, 0.205)), and 6h (t = 0.64225, df = 8, p-value = 0.5387, mean difference = 0.0478, 95% CI = (−0.124, 0.219)) time-interval treatment groups. However, the 24h time-interval treatment group showed a significant difference between basal and final heating rates for individuals (t = 2.8825, df = 9, p-value = 0.0181, mean difference = 0.1127, 95% CI = (0.024, 0.201) with final heating rates (mean = 0.87 °C/min) on average being higher than basal heating rates (mean = 0.98 °C/min).

One-way ANOVAs found no significant influence of sex, mass, SVL, or time-interval treatment group on basal heating rates across all time-interval treatment groups (p-values > 0.05) (Table 1). After considering time-interval treatment group, sex, mass measured before basal CT_max_ (or final CT_max_) experiments, and SVL showed no significant effects on final heating rates (all p-values > 0.05) except for the interaction between time-interval treatment group and mass (for mass measured before basal CT_max_ p-value =0.0423 and for mass measured before final CT_max_ experiments p-value = 0.0216) (Table 1). Cook’s Distance tests indicated that no individuals had a Cook’s Distance > 1.

**Table 1:** Influence of sex, mass, SVL, and time-interval treatment group on basal and final heating rates. Sex, mass measured before basal CT_max_ experiments, SVL, and time-interval group were tested separately on basal heating rates. Mass measured before basal and final CT_max_ experiments, SVL, and sex were tested separately with time-interval group as a covariate on final heating rates. P-values < 0.05 are in bold to represent statistical significance (df = degrees of freedom; Sum Sq = sum of squares; Mean Sq = mean square).

| Source | Df | Sum Sq | Mean Sq | F-value | p-value |
| --- | --- | --- | --- | --- | --- |
| **Basal Heating Rate** |  |  |  |  |  |
| Sex | 1 | 0.0221 | 0.02206 | 1.601 | 0.217 |
| Mass | 1 | 0.0394 | 0.03937 | 3.001 | 0.095 |
| SVL | 1 | 0.0001 | 8.4 x 10<sup>-5</sup> | 0.006 | 0.94 |
| Treatment Group | 2 | 0.0027 | 0.001349 | 0.089 | 0.915 |
| **Final Heating Rate** |  |  |  |  |  |
| Sex | 1 | 0.0010 | 0.00097 | 0.030 | 0.864 |
| Treatment Group | 2 | 0.0228 | 0.01139 | 0.354 | 0.706 |
| Treatment Group:Sex | 2 | 0.0238 | 0.01188 | 0.369 | 0.695 |
| Mass (before basal CT<sub>max</sub>) | 1 | 0.0324 | 0.03238 | 1.356 | 0.2567 |
| Treatment Group | 2 | 0.0228 | 0.01139 | 0.477 | 0.6271 |
| Treatment Group:Mass | 2 | 0.1750 | 0.08752 | 3.665 | **0.0423** |
| Mass (before final CT<sub>max</sub>) | 2 | 0.0282 | 0.01408 | 0.753 | 0.4834 |
| Treatment Group | 1 | 0.0425 | 0.04251 | 2.273 | 0.1466 |
| Treatment Group:Mass | 2 | 0.1732 | 0.08658 | 4.629 | **0.0216** |
| SVL | 1 | 0.0022 | 0.00220 | 0.073 | 0.789 |
| Treatment Group | 2 | 0.0228 | 0.03438 | 1.143 | 0.337 |
| Treatment Group:SVL | 2 | 0.0688 | 0.01138 | 0.388 | 0.682 |

### 3.3 Thermal tolerance and heat hardening capacity

Average basal CT_max_ across all individuals for all time-interval treatment groups was 41.07 °C (Table 2). Final CT_max_ averaged across individuals was highest for the 3h time-interval treatment group (mean = 41.71 °C) compared to the averages across individuals for the 6h (mean = 41.20 °C) and 24h treatment groups (mean = 40.87 °C) (Table 3; Figure 1A).

**Figure 1:**
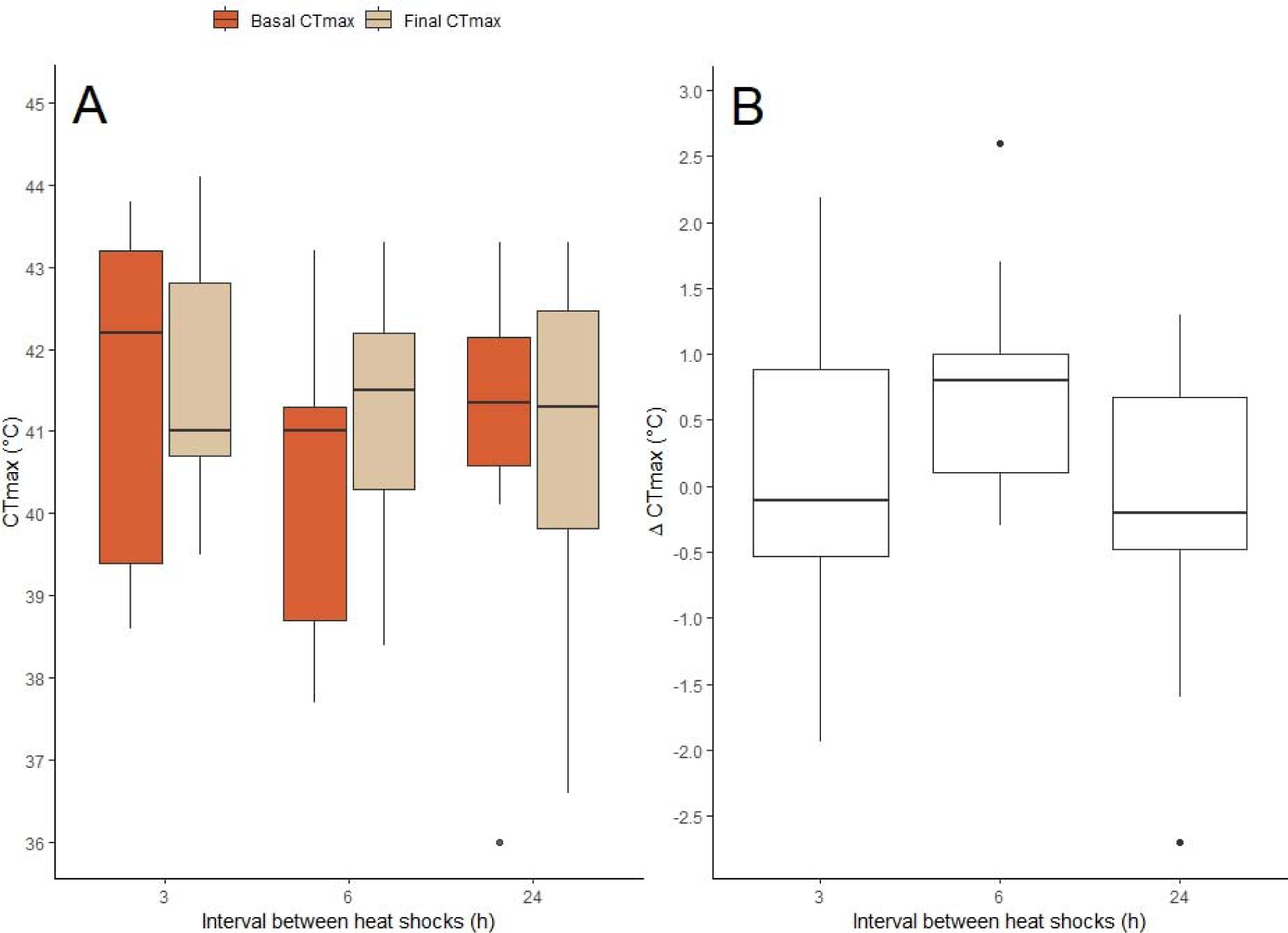
Boxplots showing (A) basal CT_max_ and final CT_max_ for all individuals of each time-interval treatment group and (B) heat hardening (change in CT_max_ between basal CT_max_ and final CT_max_) for all individuals of each time-interval treatment group.

**Table 2:** Mean and standard deviation of CT_max_ values with standard error of measurement (SEM) in parentheses of *E. macularius* before (basal CT_max,_ treatment group 0h) and after heat hardening (final CT_max,_ 3h, 6h, or 24h) for each time-interval treatment group. *n* = number of individuals.

| Treatment Group | CT<sub>max</sub> (°C) | *n* |
| --- | --- | --- |
| 0h | 41.07±1.99 (0.37) | 28 |
| 3h | 41.71±1.48 (0.49) | 9 |
| 6h | 41.20±1.68 (0.56) | 9 |
| 24h | 40.87±2.09 (0.69) | 10 |

**Table 3:** Summary of Pearson’s product-moment correlation analysis to test for the relationship between basal CT_max_ and change in CT_max_ within time-interval treatment groups (df = degrees of freedom, CI = confidence interval).

| Treatment Group | t-value | df | p-value | r | 95% CI |
| --- | --- | --- | --- | --- | --- |
| 3h | 0.297 | 7 | 0.775 | 0.111 | -0.596, 0.772 |
| 6h | -1.209 | 7 | 0.265 | -0.415 | -0.846, 0.343 |
| 24h | -0.804 | 8 | 0.444 | -0.273 | -0.770, 0.430 |

In comparison to native microclimate temperatures, basal CT_max_ averaged across individuals of *E. macularius* (mean = 41.07 °C) was found to be higher than native shade microclimate temperatures across the day and season (mean = 19.22 °C) (Supp. Materials Figure S2). Basal CT_max_ for *E. macularius* was also higher than the native sun microclimate temperatures across the day and season (mean = 22.14 °C) (Supp. Materials Figure S2). Maximum temperatures in the species’ native habitat reach 59°C and 43°C across the year in the sun and shade, respectively (Supp. Materials Figure S2).

Pearson’s product-moment analyses found no significant correlations between basal CT_max_ and change in CT_max_ (final – basal) for the 3h, 6h, or 24h time-interval treatment groups (p-values > 0.05) (Table 3). Average change in CT_max_ was found to be highest for individuals within the 6h time-interval treatment group (mean = 0.76 °C) compared to the average change in CT_max_ for individuals within the 3h (mean = 0.03 °C) and 24h (mean = −0.23 °C) treatment groups (Figure 1B).

The initial general linear model (GLM) found no effect of time interval treatment group, sex, or mass on basal CT_max_ (p-values > 0.05) (Supp. Materials Table S1a); however, SVL did show a significant effect on basal CT_max_ (p-value = 0.00359) (Supp. Materials Table S1a). This result was confirmed for the GLM with only SVL used as a variable in the model (p-value = 0.0297) (Table 4). For both sexes, individuals with higher SVL tended to have lower basal CT_max_.

**Table 4:** Summary of general linear models (GLMs) to test for the influence of significant variables based on the initial more complex GLM models (Supp. Table S1) on basal CT_max_, final CT_max_, and change in CT_max_. GLMs include mass measured before final CT_max_ experiments for final CT_max_ and change in CT_max_ models. Statistically significant p-values are shown in bold. SVL = snout-vent length (measured in cm). Mass measured in grams.

| Coefficients | Estimate | Std. Error | t-value | p-value |
| --- | --- | --- | --- | --- |
| *(a) glm(Basal $CT_{max}$ ~ SVL, family = "gaussian") AIC: 117.82* |  |  |  |  |
| (Intercept) | 56.6367 | 6.7742 | 8.361 | **7.66 x 10<sup>-9</sup>** |
| SVL (cm) | -1.3179 | 0.5728 | -2.301 | **0.0297** |
| Null deviance: 107.057 on 27 df; Residual deviance: 88.947 on 26 df |  |  |  |  |
| *(b) glm(Final $CT_{max}$ ~ Group\*Basal $CT_{max}$ + Mass, family = "gaussian") AIC: 87.621* |  |  |  |  |
| (Intercept) | 10.283 | 7.141 | 1.440 | 0.165 |
| Group (3h) | 44.562 | 10.311 | 4.322 | **0.000331** |
| Group (6h) | 4.781 | 12.745 | 0.375 | 0.711 |
| Basal $CT_{max}$ | 0.847 | 0.169 | 4.991 | **7.01 x 10<sup>-5</sup>** |
| Mass (g) | -0.089 | 0.0325 | -2.734 | **0.012790** |
| Group(3h):Basal $CT_{max}$ | -1.057 | 0.248 | -4.253 | **0.000390** |
| Group(6h):Basal $CT_{max}$ | -0.103 | 0.312 | -0.333 | 0.742 |
| Null deviance: 82.241 on 26 df; Residual deviance: 22.435 on 20 df |  |  |  |  |
| *(c) glm(Change in $CT_{max}$ ~ Mass, family = "gaussian") AIC = 82.106* |  |  |  |  |
| (Intercept) | 3.919 | 1.396 | 2.806 | **0.00957** |
| Mass (g) | -0.080 | 0.029 | -2.773 | **0.01034** |
| Null deviance: 34.637 on 26 df, Residual deviance: 26.488 on 25 df |  |  |  |  |

On the other hand, the initial GLM including all the variables indicated that final CT_max_ was significantly affected by time-interval treatment group (p-value = 0.000375), basal CT_max_ (p-value = 0.000434), and the interaction between time-interval treatment group and basal CT_max_ (p-value = 0.000440) (Supp. Materials Table S1b). These results were confirmed when the GLM was re-run using a simpler model including only significant variables based on the initial more complex model (data not shown).

Finally, change in CTmax was not affected by sex, mass measured before basal CT_max_ experiments, SVL, basal CT_max_, time-interval treatment group, or the interaction between time-interval treatment group and basal CT_max_ (p-values > 0.05) (Supp. Materials Table S1c).

Analyses repeated using the mass of all individuals measured before final CT_max_ experiments generally confirmed the results obtained using the mass measured before basal CT_max_. The only difference found was that in the case of the mass measured before final CT_max_, the mass of the individuals had an influence on final CT_max_ and on the change in CT_max_ (p-value = 0.03 in both cases, data not shown). Furthermore, we found that individuals with higher body mass exhibited lower final CT_max_ and lower change in CT_max_ (plasticity) compared to individuals with lower body mass (p-value = 0.012 and 0.010, respectively) (Table 4). Cook’s Distance tests identified no individuals with Cook’s Distance > 1. When analyses were re-run using simpler models including only significant variables (Supp. Materials Table S1), we found that across time-interval treatment groups, there is a significant positive relationship between basal CT_max_ and final CT_max_ (p-value = 0.00007) (Table 4). Linear regression models reveal that for the 6h (r^2^ = 0.7435; F-statistic = 20.29 on 1 and 7 df; p-value = 0.00278; slope = 0.7883; standard error = 0.1750; t-value = 4.504) and 24h (r^2^ = 0.6978; F-statistic = 18.47 on 1 and 8 df; p-value = 0.00262; slope = 0.8424; standard error = 0.190; t-value = 4.298) treatment groups, individuals with high basal CT_max_ tended to also have high final CT_max_. However, for the 3h treatment group, there was no significant relationship between basal CT_max_ and final CT_max_ (r^2^ = 0.1097; F-statistic = 0.8626 on 1 and 7 df; p-value = 0.38393; slope = −0.2378; standard error = 0.2560; t-value = - 0.929).

## 4. Discussion

Short-term heat hardening (thermal tolerance plasticity) has the potential to buffer the effects of increased warming caused by heatwaves on organisms, especially for species which already occupy harsh xeric habitats. In this study, we determined the basal thermal tolerance (basal CT_max_) and heat hardening capacity (change in CT_max_ between basal and final CT_max_) in one such species, the crepuscular gecko, *Eublepharis macularius*. Consistent with the known male-biased sexual size dimorphism in native populations of *E. macularius* (Kratochvíl & Frynta, 2002), sexual-size dimorphism was shown between males and females studied in our work, with males having larger SVL on average than females.

To our knowledge, in this study, we estimated for the first time the influence of variables on heating rates, the rates at which the organism increases its body temperature per minute, in heat hardening studies. While other studies have looked at the influence of heating rates for CT_max_ studies, this approach is not normally included in studies on heat hardening. We found that basal and final heating rates for CT_max_ experiments did not differ across and within time-interval treatment groups, except for the 24h time-interval treatment group, which showed a mean difference = 0.1127 °C/min between basal and final heating rates. We also found that sex, mass, and SVL did not influence basal and final heating rates except for an interaction between mass and time-interval group for final heating rates. To note that our data indicate that the individual mass does not differ among the different treatment groups. These results support the comparison of basal and final CT_max_ data to study heat hardening in this species as they validate the consistency of heating rates for measuring CT_max_ regardless of the sex, mass, SVL and time-interval treatment group. We suggest that future research incorporate analyses of the heating rates used for measurements of CT_max_, in order to take into account potential variation in basal and final CT_max_ due to different heating rates.

We found that heat hardening capacity - the aptitude of an organism to plastically adjust its thermal limit - in *E. macularius* was not affected by the basal CT_max_ of the individual. Thus, we reject the TOH - according to which heat hardening is higher in organisms with lower basal thermal tolerance and vice-versa - for *E. macularius*. In fact, we found that individuals with relatively low basal CT_max_ did not exhibit greater change in CT_max_ and vice versa across the 3h and 6h time-interval treatment groups. Although comparative studies of heat hardening capacity across squamates are lacking, heat hardening capacity in different lizard species have generally rejected the TOH with the exception of two tropical species, *Lampropholis coggeri* (Phillips et al., 2016; Gunderson, 2023) and *Hemidactylus frenatus* (Lapwong et al., 2021). Furthermore, the TOH also vary in its support across other taxa of ectotherms. For example, one species of crustaceans has shown strong support for the TOH across multiple studies, whereas two fish species have shown no support for the TOH (Gunderson, 2023). Differences in the trade-off between thermal tolerance and plasticity across and within species could be caused by different mechanisms of selection (i.e., correlational, opposing) for these two traits based on the organism’s native environment (van Heerwaarden & Kellermann, 2020). For example, high or low thermal tolerance together with high or low plasticity may be adaptive in some environments, whereas only high plasticity or only high thermal tolerance may be adaptive in others (van Heerwaarden & Kellermann, 2020) because of differences in the species thermal environment. Uncovering how widespread TOH may be and which factors may influence it will require more research, since heat hardening has been studied in a limited number of species.

Basal CT_max_ measured in individuals of *E. macularius* was found to be relatively lower (mean = 41.07 °C) compared to other species within the family *Eublepharidae* (mean across species = 42.3 °C; Clusella-Trullas & Chown, 2014); however, thermal tolerance has been scarcely studied within *Eublepharidae*. The two species that have been studied, *Coleonyx brevis* (CT_max_ = 41.6 °C; SVL = 5.1 cm) and *Coleonyx variegatus* (CT_max_ = 43.0 °C; SVL = 6.5 cm) (Dial & Grismer, 1992), have much smaller body size than *E. macularius*, which could explain the higher thermal tolerance exhibited within these two closely related species. To further corroborate the relationship between body size and thermal tolerance observed in ectotherms (Peralta-Maraver & Rezende, 2021), our data indicate that in our dataset, independently of the sex of the individuals, individuals with smaller body size (SVL) were found to have higher basal CT_max_ compared to larger individuals. Intraspecific differences in body size have been shown to influence CT_max_ measurements in other lizard species (Brusch IV, 2016; Claunch, 2021). Larger individuals tend to have lower CT_max_ in four *Sceloporus* species, which has been hypothesized to be affected by thermal inertia, metabolic/oxygen delivery constraints, or a combination of both (Claunch et al, 2021). Smaller ectotherms have also been shown to have higher thermal tolerance in response to acute heat stress compared to larger ectotherms (Peralta-Maraver & Rezende, 2021). The higher basal thermal tolerance of *E. macularius* in our study (mean ∼ 41°C) than the mean microclimate temperatures observed in its native environment (mean sun ∼ 22°C; mean shade ∼ 19 °C) suggests that although this species is primarily active at dawn and dusk and hibernates during the winter months of the year (Rawat et al., 2019), it has adapted its upper thermal tolerance to the highest possible temperatures experienced in its native environment across the day and season. Although, maximum temperatures in the species’ native habitat reach 59°C and 43°C across the year in the sun and shade, respectively, the mean temperature is close to 40°C during the mid-day hours in the sun across the year and above 25°C for several months during the year (Supp. Materials Figure S2).

Final CT_max_ – the maximum temperature that the organism could tolerate after 3, 6, or 24 hours - was found to be influenced by both basal CT_max_ and time-interval treatment group for individuals of *E. macularius*. Individuals within the 6h and 24h treatment groups showed a significant positive relationship between basal and final CT_max_ in which individuals with relatively low or high basal CT_max_ also showed relatively low or high final CT_max_ after 6 hours or 24 hours from initial heat shock, respectively. In the only other species of geckos studied so far, *Hemidactylus frenatus*, heat hardening experiments also found that basal CT_max_ also significantly affected final CT_max_ (Lapwong et al., 2021). To note that in our study, individuals within the 3h time-interval group did not show a significant relationship between basal and final CT_max_. These results suggest that individuals of *E. macularius* may require a certain amount of time in between heat shocks (i.e, greater than 3 hours) to physiologically recover from initial heat stress and increase their thermal tolerance through the upregulation of heat shock proteins at the cellular level in order to prepare for the next heat stress event (Angilletta et al., 2009; Gangloff & Telemeco 2018). Furthermore, individuals of *E. macularius*, regardless of sex, showed a significant negative relationship between body mass and final CT_max_ where individuals with higher body mass exhibited a lower final CT_max_, similar to what observed for basal CT_max_ and body size (SVL) in our experiments. Thus, both body mass and size can impact thermal tolerance for individuals of *E. macularius*, which is consistent with the previously discussed trend found not only across lizard species (Brusch IV, 2016; Claunch, 2021), but also across ectotherms (Peralta-Maraver & Rezende, 2021).

Our results also support our hypothesis that changes in CT_max_ (heat hardening capacity) would be highest after 6 hours since initial heat shock for individuals of *E. macularius*, in agreement with what observed in other lizards (Phillips et al., 2016; Lapwong et al., 2021; Gilbert & Miles, 2019). In addition, the time course of heat hardening response across 24 hours for individuals of *E. macularius* is very similar to what found in other lizard species, where the magnitude of plasticity (i.e., change in thermal tolerance) increases to a set point and then decreases close to zero across 24 hours. To note that although this trend has been described for some diurnal species (e.g., *Anolis carolinensis* and *Lampropholis coggeri*; Deery et al., 2021, Phillips et al., 2016), it was not observed in the diurnal lizard *Anolis sagrei* (Deery et al., 2021) or in the crepuscular gecko *Hemidactylus frenatus* (Lapwong et al., 2021), in which thermal tolerance plasticity did not increase over time. Thus, the ability to respond quickly to heat stress by plasticly adjusting thermal tolerance may vary across and within species independently of the time of activity and may be instead more related to the seasonal variation and predictability of thermal variation for native environmental temperatures (e.g., Phillips et al., 2016; Deery et al., 2021). To highlight that the individuals of *E. macularius* used in this study are of captive origin and as such thermal tolerance plasticity may be lower than what it would be observed on wild individuals, since the native thermal environment of this species varies considerably across the day and season (Supp. Materials Figure S2).

Finally, although we found an influence of SVL and body mass on basal and final CT_max_, respectively, we found that changes in CT_max_ (heat hardening capacity) were not influenced by body size. On the other hand, changes in CT_max_ were influenced by body mass in the studied species, with individuals with lower body mass experiencing greater change in CT_max_ compared to individuals with higher body mass. Acclimation capacity has been shown to be positively influenced by organism body size and mass across ectotherm species (Rohr et al., 2018; Brown et al., 2004; Kingsolver & Huey, 2008; Pörtner et al., 2017). Larger individuals (or species) have been found to have greater acclimation capacity (i.e., change in the upper thermal tolerance relative to change in mean temperature, Claussen, 1977), although smaller organisms have a faster acclimation rate (Rohr et al., 2018). As a consequence, during short-term acclimation, smaller organisms may show greater plasticity due to their faster acclimation (Rohr et al., 2018). Thus, although our results where smaller individuals have higher change in CT_max_ (heat hardening/plasticity) seem to not support the general relationship between body mass and acclimation capacity (heat hardening) as observed in other species, this may be due to individuals being exposed to short-term instead of long-term thermal stress. In addition, smaller organisms have higher mass-specific metabolic rates and lower oxygen demand and delivery constraints compared to organisms with greater body mass, which has been theorized to aid in the faster acclimation responses of smaller organisms (Brown et al., 2004; Kingsolver & Huey, 2008; Pörtner et al., 2017)

## 5. Conclusions

Our results suggest that basal thermal tolerance does not influence heat hardening capacity in *E. macularius*, indicating that individuals with lower basal thermal tolerance do not compensate for this by possessing a greater capacity to plasticly adjust their upper thermal limit. Although this may be the results of an adaptation to captivity conditions, where the developmental environment may be more controlled and homogeneous than in the wild, the lack of trade-off between basal thermal tolerance and plasticity seems to largely vary within and across different species. To further understand the interplay between basal thermal tolerance and plasticity in organisms, future studies should focus not only on collecting similar data across species and populations of ectotherms, but also on investigating the mechanisms underlying differences in selection influencing this trade-off. Additionally, our data support a strong similitude in heat hardening capacity between the studied captive population of *E. macularius* and what observed in other squamates sampled from the wild. These results suggest that although plasticity of thermal tolerance via heat hardening may be somehow influenced due to captivity, the species’ biology, physiology, and evolutionary history may have a stronger impact. Our work, together with the work done so far on thermal tolerance plasticity in ectotherms, highlights the need of carrying out comparative studies across species and collecting data across different levels of complexity, from molecular to ecological and physiological, to uncover the abiotic factors influencing thermal tolerance plasticity and the molecular mechanisms underlying the observed variation.

Finally, heat stress caused by climate change has been predicted to negatively impact ectothermic species, especially for tropical and mid-latitude organisms (Kingsolver et al., 2013). This study increases our understanding of how quickly organisms can respond to daily thermal extremes. To note that the studies available so far on this topic do not address responses across multiple or prolonged heat stress events. As heat waves and heat stress are predicted to increase in length and frequency in the future (Stillman et al., 2019; Angélil et al., 2017; Guo et al., 2018; Pachauri et al., 2014), to properly assess how organisms will be impacted, it will be necessary for future studies to also investigate variation in thermal tolerance plasticity across days or weeks.

Taken all together, our study contributes to the limited data available so far on thermal tolerance plasticity in squamates, focusing on an extremely understudied group for this topic (geckos), a species mostly active during crepuscular/early night hours, and a population that has been captive bred for several (unknown) generations. Furthermore, our study also provides methodological guidelines on checking potential variation (in heating rates) that may influence results and their interpretation of studies on thermal tolerance plasticity, independently of the studied species.

## Acknowledgements

We are thankful to Alex Gunderson, Vincent Farallo, Miguel Carretero, Catarina Rato, and Eric Gangloff for helpful advice with designing and setting up experiments, and to Patrick Gillevet for technical help with the temperature-controlled room used for the experiments. We are thankful to Patrick Gillevet and Rebecca Forkner for providing helpful comments on an earlier version of this manuscript.

## Author Contributions

Conceptualization: E.W., Y.C.; Methodology: E.W.; Y.C.; Formal Analysis: E.W.; Resources: Y.C.; Investigation: E.W., S.K., G.W.; Writing - original draft: E.W., Y.C.; Writing - review & editing: S.K., G.W.; Supervision: Y.C.; Project administration: Y.C.

## Data accessibility

Full dataset and Rscript used for analyses will be available on Dryad *after manuscript acceptance*.

## References

Alexander, L. V., Zhang, X., Peterson, T. C., Caesar, J., Gleason, B., Klein Tank, A. M. G., … & Vazquez_JAguirre, J. L. (2006). Global observed changes in daily climate extremes of temperature and precipitation. Journal of Geophysical Research: Atmospheres, 111(D5).

Angélil, O., Stone, D., Wehner, M., Paciorek, C. J., Krishnan, H., & Collins, W. (2017). An independent assessment of anthropogenic attribution statements for recent extreme temperature and rainfall events. Journal of Climate, 30(1), 5–16.

Angilletta, M. J. (2009). Thermal adaptation: a theoretical and empirical synthesis. Oxford University Press.

Bilyk, K. T., Evans, C. W., & DeVries, A. L. (2012). Heat hardening in Antarctic notothenioid fishes. Polar biology, 35(9), 1447–1451.

Bowler, K. (2005). Acclimation, heat shock and hardening. Journal of Thermal Biology, 30(2), 125–130.

Brown, J. H., Gillooly, J. F., Allen, A. P., Savage, V. M., & West, G. B. (2004). Toward a metabolic theory of ecology. Ecology, 85(7), 1771–1789.

Brusch IV, G. A., Taylor, E. N., & Whitfield, S. M. (2016). Turn up the heat: thermal tolerances of lizards at La Selva, Costa Rica. Oecologia, 180(2), 325–334.

Buckley, B. A., & Hofmann, G. E. (2002). Thermal acclimation changes DNA-binding activity of heat shock factor 1 (HSF1) in the goby Gillichthys mirabilis: implications for plasticity in the heat-shock response in natural populations. Journal of Experimental Biology, 205(20), 3231–3240.

Camacho, A., & Rusch, T. W. (2017). Methods and pitfalls of measuring thermal preference and tolerance in lizards. Journal of Thermal Biology, 68, 63–72.

Chamberlain, S., Ram, K., Barve, V., Mcglinn, D., & Chamberlain, M. S. (2017). Package ‘rgbif’. Interface to the Global Biodiversity Information Facility ‘API, 5(0.9).

Claunch, N. M., Nix, E., Royal, A. E., Burgos, L. P., Corn, M., DuBois, P. M., … & Taylor, E. N. (2021). Body size impacts critical thermal maximum measurements in lizards. Journal of Experimental Zoology Part A: Ecological and Integrative Physiology, 335(1), 96–107.

Claussen, D. L. (1977). Thermal acclimation in ambystomatid salamanders. Comparative Biochemistry and Physiology A, 58, 333–340

Clusella-Trullas, S., & Chown, S. L. (2014). Lizard thermal trait variation at multiple scales: a review. Journal of Comparative Physiology B, 184, 5–21.

Coffel, E. D., Horton, R. M., & De Sherbinin, A. (2017). Temperature and humidity based projections of a rapid rise in global heat stress exposure during the 21st century. Environmental Research Letters, 13(1), 014001.

Cook, R. D. (1977). Detection of influential observation in linear regression. Technometrics, 19(1), 15–18.

Daugaard, M., Rohde, M., & Jäättelä, M. (2007). The heat shock protein 70 family: Highly homologous proteins with overlapping and distinct functions. FEBS letters, 581(19), 3702–3710.

Dayananda, B., Gray, S., Pike, D., & Webb, J. K. (2016). Communal nesting under climate change: fitness consequences of higher incubation temperatures for a nocturnal lizard. Global Change Biology, 22(7), 2405–2414.

Deery, Rej, J. E., Haro, D., & Gunderson, A. R. (2021). Heat hardening in a pair of Anolis lizards: constraints, dynamics and ecological consequences. Journal of Experimental Biology, 224(Pt 7).

Dial, B. E., & Grismer, L. L. (1992). A phylogenetic analysis of physiological-ecological character evolution in the lizard genus Coleonyx and its implications for historical biogeographic reconstruction. Systematic Biology, 41(2), 178–195.

Easterling, D. R., Horton, B., Jones, P. D., Peterson, T. C., Karl, T. R., Parker, D. E., … & Folland, C. K. (1997). Maximum and minimum temperature trends for the globe. Science, 277(5324), 364–367.

Froelicher, T., Fischer, E. M., Gruber, N., Striegel, S., & Laufkötter, C. (2019, January). Marine Heat Waves under Global Warming (Invited Presentation). In 99th American Meteorological Society Annual Meeting. AMS.

Gamble, T., Greenbaum, E., Jackman, T. R., & Bauer, A. M. (2015). Into the light: diurnality has evolved multiple times in geckos. Biological Journal of the Linnean Society, 115(4), 896–910.

Gangloff, E. J., & Telemeco, R. S. (2018). High temperature, oxygen, and performance: Insights from reptiles and amphibians. Integrative and Comparative Biology, 58(1), 9–24.

Gilbert, A. L., & Miles, D. B. (2019). Antagonistic responses of exposure to sublethal temperatures: adaptive phenotypic plasticity coincides with a reduction in organismal performance. The American Naturalist, 194(3), 344–355.

Glimm, T., Kiskowski, M., Moreno, N., & Chiari, Y. (2021). Capturing and analyzing pattern diversity: an example using the melanistic spotted patterns of leopard geckos. PeerJ, 9, e11829.

Grismer, L. L., Wood Jr, P. L., Ngo, V. T., & Murdoch, M. L. (2015). The systematics and independent evolution of cave ecomorphology in distantly related clades of Bent-toed Geckos (Genus Cyrtodactylus Gray, 1827) from the Mekong Delta and islands in the Gulf of Thailand. Zootaxa, 3980(1), 106–126.

Gunderson, A. R. (2023). Trade_Joffs between baseline thermal tolerance and thermal tolerance plasticity are much less common than it appears. Global Change Biology.

Gunderson, A. R., Dillon, M. E., & Stillman, J. H. (2017). Estimating the benefits of plasticity in ectotherm heat tolerance under natural thermal variability. Functional Ecology, 31(8), 1529–1539.

Gunderson, A. R., & Stillman, J. H. (2015). Plasticity in thermal tolerance has limited potential to buffer ectotherms from global warming. Proceedings of the Royal Society B: Biological Sciences, 282(1808), 20150401.

Guo, Y., Gasparrini, A., Li, S., Sera, F., Vicedo-Cabrera, A. M., de Sousa Zanotti Stagliorio Coelho, M., … & Tong, S. (2018). Quantifying excess deaths related to heatwaves under climate change scenarios: A multicountry time series modelling study. PLoS medicine, 15(7), e1002629.

Hamdoun, A. M., Cheney, D. P., & Cherr, G. N. (2003). Phenotypic plasticity of HSP70 and HSP70 gene expression in the Pacific oyster (Crassostrea gigas): implications for thermal limits and induction of thermal tolerance. The Biological Bulletin, 205(2), 160–169.

Hastings, B. T., Melnyk, A., Ghyabi, M., White, E., Barroso, F. M., Carretero, M. A., … & Chiari, Y. (2023). On the role of melanistic coloration on thermoregulation in the crepuscular gecko Eublepharis macularius. bioRxiv, 2023−05.

Heinicke, M. P., Jackman, T. R., & Bauer, A. M. (2017). The measure of success: geographic isolation promotes diversification in Pachydactylus geckos. BMC Evolutionary Biology, 17(1), 1–17.

Hoffmann, A. A., Chown, S. L., & Clusella_JTrullas, S. (2013). Upper thermal limits in terrestrial ectotherms: how constrained are they?. Functional Ecology, 27(4), 934–949.

Horowitz, M. (2001). Heat acclimation: phenotypic plasticity and cues to the underlying molecular mechanisms. Journal of Thermal Biology, 26(4-5), 357–363.

Hu, J. T., Chen, B., & Li, Z. H. (2014). Thermal plasticity is related to the hardening response of heat shock protein expression in two Bactrocera fruit flies. Journal of Insect Physiology, 67, 105–113.

Huey, R. B. (1991). Physiological consequences of habitat selection. The American Naturalist, 137, S91–S115.

Janzen, D. H. (1967). Why mountain passes are higher in the tropics. The American Naturalist, 101(919), 233–249.

Karl, T. R., Kukla, G., Razuvayev, V. N., Changery, M. J., Quayle, R. G., Heim Jr, R. R., … & Fu, C. B. (1991). Global warming: Evidence for asymmetric diurnal temperature change. Geophysical Research Letters, 18(12), 2253–2256.

Katlein, N., Ray, M., Wilkinson, A., Claude, J., Kiskowski, M., Wang, B., Glaberman, S. & Chiari, Y. (2022). Does colour impact responses to images in geckos?. Journal of Zoology, 317(2), 138–146.

Kearney, M. R., & Porter, W. P. (2017). NicheMapR - an R package for biophysical modeling: The microclimate model. Ecography, 40(5), 664–674.

Kelly, C., & Price, T. D. (2005). Correcting for regression to the mean in behavior and ecology. The American Naturalist, 166(6), 700–707.

Khan, M. S. (2009). Natural history and biology of hobbyist choice leopard gecko Eublepharis macularius. *Talim ul Islam College, Rabwah*, Pakistan.

Khan, N., Shahid, S., Ahmed, K., Wang, X., Ali, R., Ismail, T., & Nawaz, N. (2020). Selection of GCMs for the projection of spatial distribution of heat waves in Pakistan. Atmospheric Research, 233, 104688.

Kingsolver, J., & Huey, R. (2008). Size, temperature, and fitness: three rules. Evolutionary Ecology Research, 10(2), 251–268.

Kingsolver, J. G., Diamond, S. E., & Buckley, L. B. (2013). Heat stress and the fitness consequences of climate change for terrestrial ectotherms. Functional Ecology, 27(6), 1415–1423.

Kiskowski, M., Glimm, T., Moreno, N., Gamble, T., & Chiari, Y. (2019). Isolating and quantifying the role of developmental noise in generating phenotypic variation. PLOS Computational Biology, 15(4), e1006943.

Kratochvíl, L., & Frynta, D. (2002). Body size, male combat and the evolution of sexual dimorphism in eublepharid geckos (Squamata: Eublepharidae). Biological Journal of the Linnean Society, 76(2), 303–314.

Kregel, K. C. (2002). Invited review: heat shock proteins: modifying factors in physiological stress responses and acquired thermotolerance. Journal of Applied Physiology, 92(5), 2177–2186.

Lapwong, Y., Dejtaradol, A., & Webb, J. K. (2021). Plasticity in thermal hardening of the invasive Asian house gecko. Evolutionary Ecology, 35(4), 631–641.

Madeira, D., Narciso, L., Cabral, H. N., & Vinagre, C. (2012). Thermal tolerance and potential impacts of climate change on coastal and estuarine organisms. Journal of Sea Research, 70, 32–41.

Madeira, D., Narciso, L., Cabral, H. N., Diniz, M. S., & Vinagre, C. (2014). Role of thermal niche in the cellular response to thermal stress: Lipid peroxidation and HSP70 expression in coastal crabs. Ecological indicators, 36, 601–606.

Manenti, T., Sørensen, J. G., Moghadam, N. N., & Loeschcke, V. (2014). Predictability rather than amplitude of temperature fluctuations determines stress resistance in a natural population of Drosophila simulans. Journal of Evolutionary Biology, 27(10), 2113–2122.

Meiri, S. (2019). What geckos are–an ecological-biogeographic perspective. Israel Journal of Ecology and Evolution, 66(3−4), 253–263.

Morgan, R., Finnøen, M. H., & Jutfelt, F. (2018). CTmax is repeatable and doesn’t reduce growth in zebrafish. Scientific reports, 8(1), 7099.

Mottola, G., Lopez, M. E., Vasemägi, A., Nikinmaa, M., & Anttila, K. (2022). Are you ready for the heat? Phenotypic plasticity versus adaptation of heat tolerance in three_Jspined stickleback. Ecosphere, 13(4), e4015.

Noer, N. K., Ørsted, M., Schiffer, M., Hoffmann, A. A., Bahrndorff, S., & Kristensen, T. N. (2022). Into the wild—a field study on the evolutionary and ecological importance of thermal plasticity in ectotherms across temperate and tropical regions. Philosophical Transactions of the Royal Society B, 377(1846), 20210004.

Oliver, P. M., Ashman, L. G., Bank, S., Laver, R. J., Pratt, R. C., Tedeschi, L. G., & Moritz, C. C. (2019). On and off the rocks: persistence and ecological diversification in a tropical Australian lizard radiation. BMC Evolutionary Biology, 19, 1–15.

Pachauri, R. K., Allen, M. R., Barros, V. R., Broome, J., Cramer, W., Christ, R., … & van Ypserle, J. P. (2014). Climate change 2014: Synthesis report. Contribution of Working Groups I, II and III to the Fifth Assessment Report of the Intergovernmental Panel on Climate Change (p. 151). IPCC.

Papenfuss, T., Shafiei Bafti, S. & Sharifi, M. 2021. Eublepharis macularius. The IUCN Red List of Threatened Species 2021: e.T164745A1072324. 10.2305/IUCN.UK.2021-3.RLTS.T164745A1072324.en. Accessed on 19 April 2022.

Peralta-Maraver, I., & Rezende, E. L. (2021). Heat tolerance in ectotherms scales predictably with body size. Nature Climate Change, 11(1), 58–63.

Phillips, Muñoz, M. M., Hatcher, A., Macdonald, S. L., Llewelyn, J., Lucy, V., & Moritz, C. (2016). Heat hardening in a tropical lizard: geographic variation explained by the predictability and variance in environmental temperatures. Functional Ecology, 30(7), 1161–1168.

Pinto, B. J., Gamble, T., Smith, C. H., Keating, S. E., Havird, J. C., & Chiari, Y. (2023). The revised reference genome of the leopard gecko (Eublepharis macularius) provides insight into the considerations of genome phasing and assembly. bioRxiv, 2023–01.

Pörtner, H. O., Bock, C., & Mark, F. C. (2017). Oxygen-and capacity-limited thermal tolerance: bridging ecology and physiology. Journal of Experimental Biology, 220(15), 2685–2696.

Rawat, Y. B., Thapa, K. B., Bhattarai, S., & Shah, K. B. (2019). First Records of the Common Leopard Gecko, Eublepharis macularius (Blyth 1854) (Eublepharidae), in Nepal. Reptiles & Amphibians, 26(1), 58–61.

Rezende, E. L., Tejedo, M., & Santos, M. (2011). Estimating the adaptive potential of critical thermal limits: methodological problems and evolutionary implications. Functional Ecology, 25(1), 111–121.

Ribeiro, P. L., Camacho, A., & Navas, C. A. (2012). Considerations for assessing maximum critical temperatures in small ectothermic animals: insights from leaf-cutting ants. PLoS One, 7(2), e32083.

Rohini, P., Rajeevan, M., & Mukhopadhay, P. (2019). Future projections of heat waves over India from CMIP5 models. Climate Dynamics, 53(1), 975–988.

Rohr, J. R., Civitello, D. J., Cohen, J. M., Roznik, E. A., Sinervo, B., & Dell, A. I. (2018). The complex drivers of thermal acclimation and breadth in ectotherms. Ecology Letters, 21(9), 1425–1439.

Ruibal, R. (1961). Thermal relations of five species of tropical lizards. Evolution, 15(1), 98–111.

Sørensen, J. G., Kristensen, T. N., & Loeschcke, V. (2003). The evolutionary and ecological role of heat shock proteins. Ecology Letters, 6(11), 1025–1037.

Rutschmann, A., Perry, C., Le Galliard, J. F., Dupoué, A., Lourdais, O., Guillon, M., Brusch, G., 4th, Cote, J., Richard, M., Clobert, J., & Miles, D. B. (2023). Ecological responses of squamate reptiles to nocturnal warming. Biological reviews of the Cambridge Philosophical Society, 10.1111/brv.13037.

Sasaki, M. C., & Dam, H. G. (2019). Integrating patterns of thermal tolerance and phenotypic plasticity with population genetics to improve understanding of vulnerability to warming in a widespread copepod. Global change biology, 25(12), 4147–4164.

Sørensen, M. H., Kristensen, T. N., Lauritzen, J. M. S., Noer, N. K., Høye, T. T., & Bahrndorff, S. (2019). Rapid induction of the heat hardening response in an Arctic insect. Biology Letters, 15(10), 20190613.

Stillman, J. H. (2019). Heat waves, the new normal: summertime temperature extremes will impact animals, ecosystems, and human communities. Physiology, 34(2), 86–100.

Telemeco, R. S., & Gangloff, E. J. (2021). Introduction to the special issue–Beyond CTMAX and CTMIN: Advances in studying the thermal limits of reptiles and amphibians. Journal of Experimental Zoology Part A: Ecological and Integrative Physiology, 335(1), 5–12.

Terblanche, J. S., Deere, J. A., Clusella-Trullas, S., Janion, C., & Chown, S. L. (2007). Critical thermal limits depend on methodological context. Proceedings of the Royal Society B: Biological Sciences, 274(1628), 2935–2943.

Tewksbury, J. J., Huey, R. B., & Deutsch, C. A. (2008). Putting the heat on tropical animals. M Science, 320(5881), 1296–1297.

Uetz, P., Slavenko, A., Meiri, S., & Heinicke, M. (2020). Gecko diversity: a history of global discovery. Israel Journal of Ecology and Evolution, 66(3−4), 117–125.

Uetz, P., Freed, P., Aguilar, R., & Hošek, J. (2023). The Reptile Database. http://www.reptile-database.org, accessed 2023-12-08.

Ulmasov, K. A., Shammakov, S., Karaev, K., & Evgen’ev, M. B. (1992). Heat shock proteins and thermoresistance in lizards. Proceedings of the National Academy of Sciences, 89(5), 1666–1670.

van Heerwaarden, B., Lee, R. F. H., Overgaard, J., & Sgrò, C. M. (2014). No patterns in thermal plasticity along a latitudinal gradient in Drosophila simulans from eastern Australia. Journal of Evolutionary Biology, 27(11), 2541–2553.

van Heerwaarden, B., & Kellermann, V. (2020). Does plasticity trade off with basal heat tolerance?. Trends in Ecology & Evolution, 35(10), 874–885.

Vose, R. S., Easterling, D. R., & Gleason, B. (2005). Maximum and minimum temperature trends for the globe: An update through 2004. Geophysical Research Letters, 32(23).

Williams, C. M., Buckley, L. B., Sheldon, K. S., Vickers, M., Pörtner, H. O., Dowd, W. W., … & Stillman, J. H. (2016). Biological impacts of thermal extremes: mechanisms and costs of functional responses matter. Integrative and Comparative Biology, 56(1), 73–84.

Zatsepina, O. G., Ulmasov, K. A., Beresten, S. F., Molodtsov, V. B., Rybtsov, S. A., & Evgen’Ev, M. B. (2000). Thermotolerant desert lizards characteristically differ in terms of heat-shock system regulation. Journal of Experimental Biology, 203(6), 1017–1025.

Zizka, A., Silvestro, D., Andermann, T., Azevedo, J., Duarte Ritter, C., Edler, D., … & Antonelli, A. (2019). CoordinateCleaner: Standardized cleaning of occurrence records from biological collection databases. Methods in Ecology and Evolution, 10(5), 744–751.

