## Supplementary Materials for "Thermal Tolerance Plasticity and Dynamics of Thermal Tolerance in *Eublepharis macularius*: Implications for Future Climate-Driven Heat Stress"

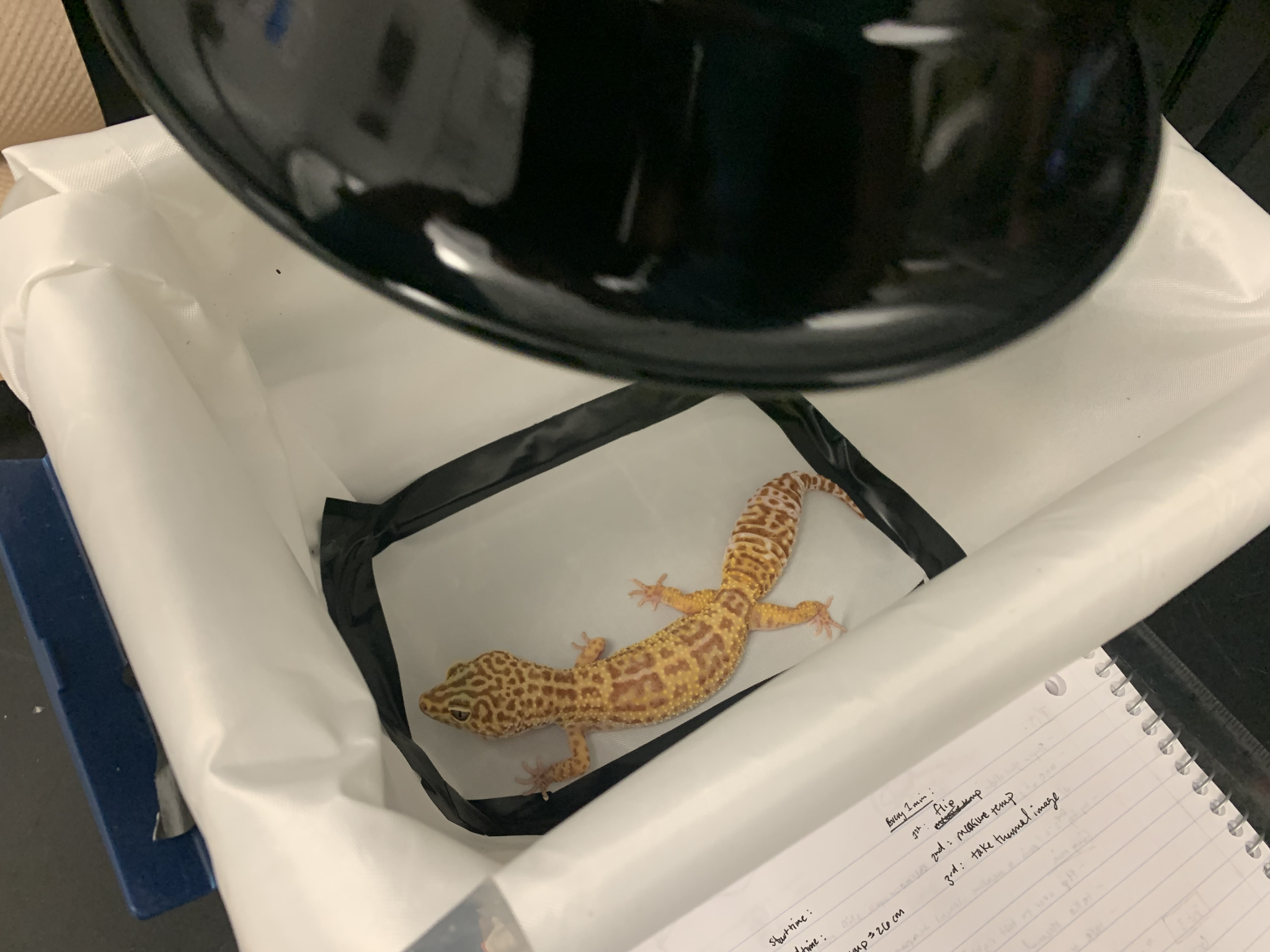

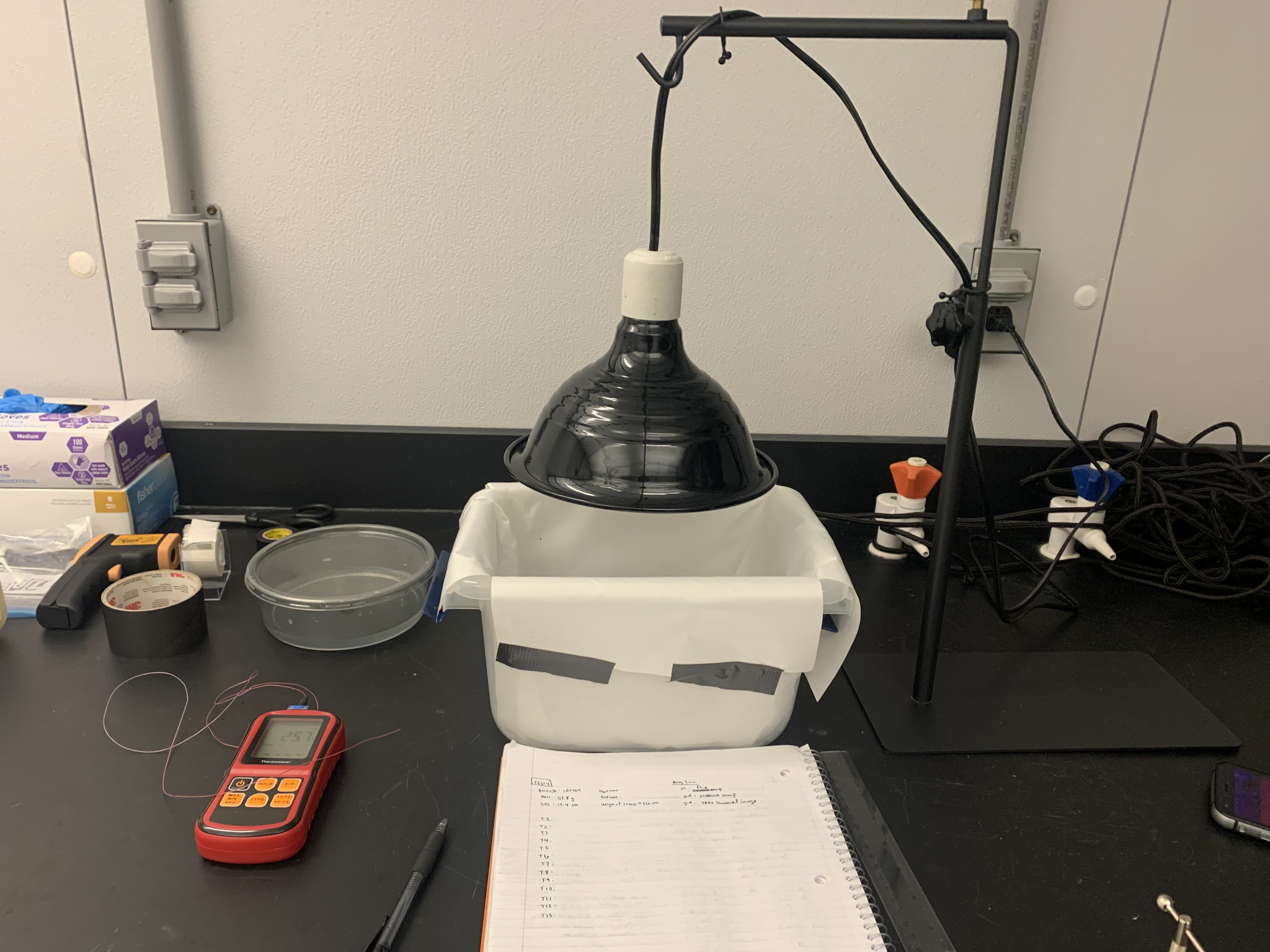

**Figure S1:** Leopard gecko placed in a plastic holding container for CT_max_ experiments (left). The experimental setup for CT_max_ experiments is shown on the right.

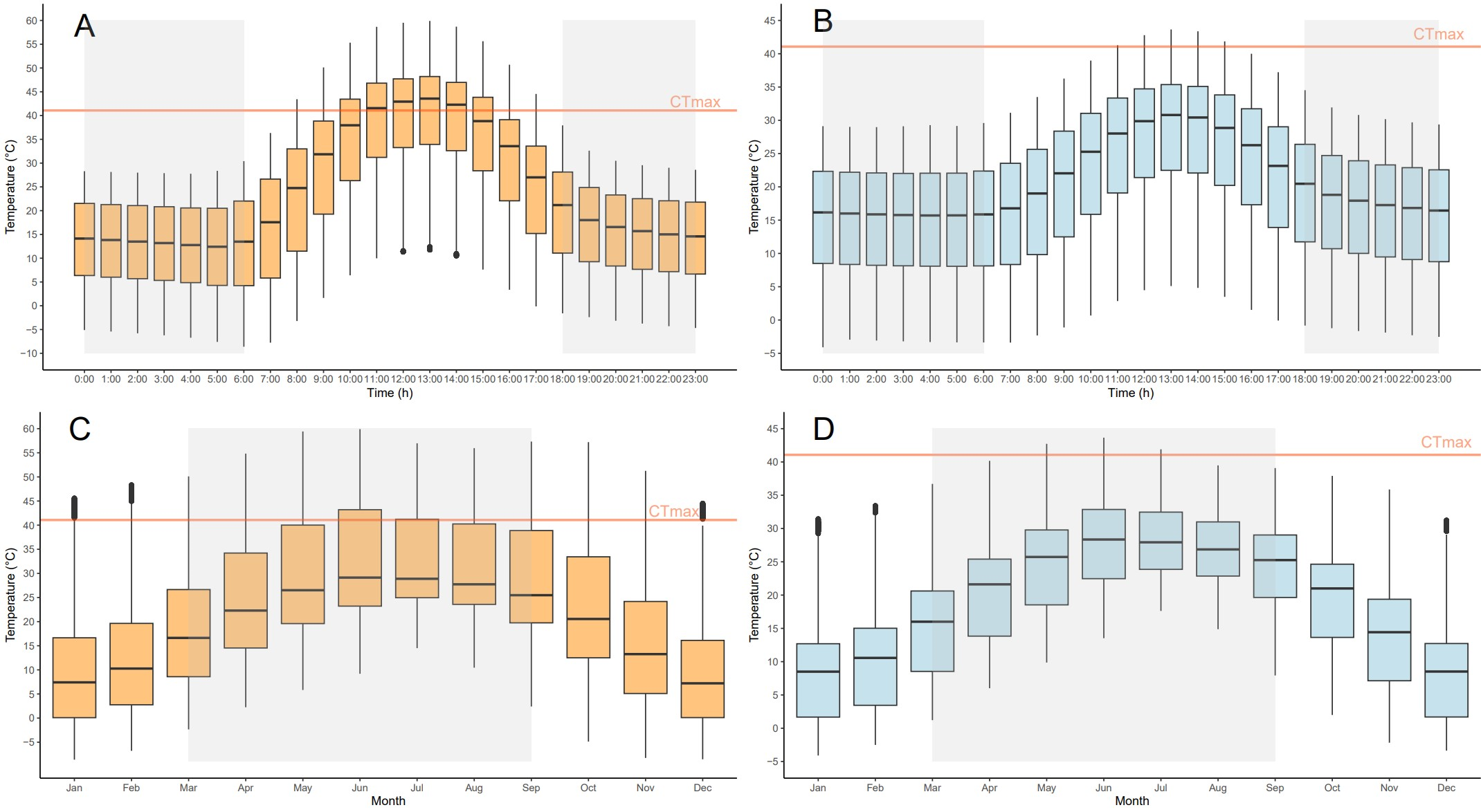

**Figure S2:** Native microclimate temperatures estimated from Global Biodiversity Information Facility (GBIF) occurrences for *E. macularius* across each hour of day for all days of the year in sun (A) and shade (B) and for each month for all days of the year in sun (C) and shade (D). Shaded boxes represent times of activity for *E. macularius* across the day (A and B) and times where hibernation does not occur across the season (C and D). Orange lines represent the mean basal CT_max_ (mean = 41.07 ^o^C) for all individuals of *E. macularius* from this study.

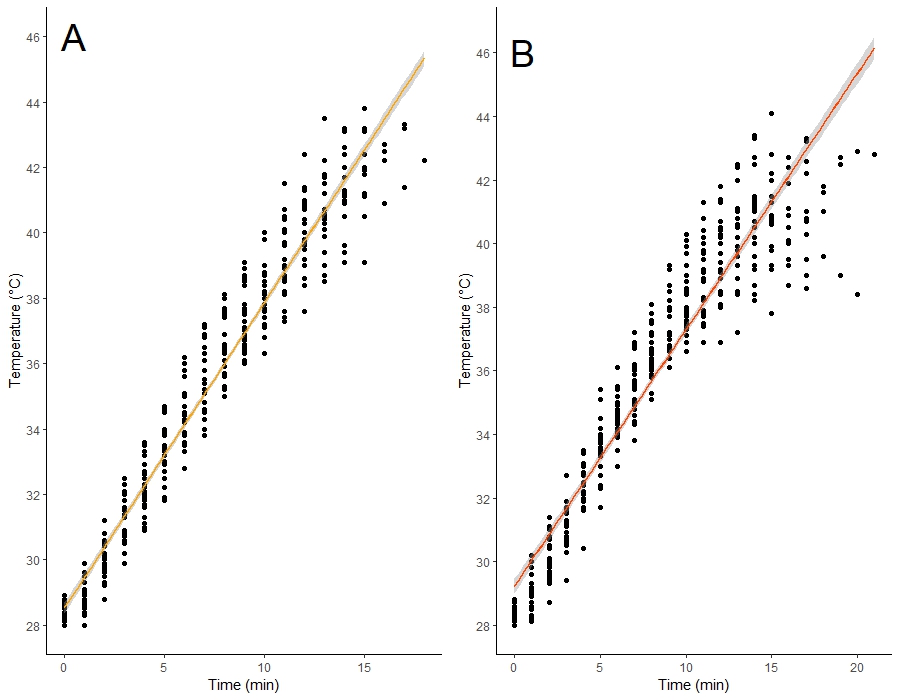

**Figure S3:** Cloacal temperatures for individual of *E. macularius* over time for basal CT_max_ experiments (A) and final CT_max_ experiments (B).

**Table S1:** Summary of general linear models (GLMs) to test for the influence of time interval treatment groups on basal CT_max_, final CT_max_, and change in CT_max_. Mass values used in each GLM included measurements taken before basal CT_max_ experiments for all individuals. Statistically significant p-values are shown in bold. SVL = snout-vent length (measured in cm). Mass measured in grams. Analyses repeated using the mass of all individuals measured before final CT_max_ experiments confirmed the results obtained using the mass measured before basal CT_max_, with the only difference that in this case, the mass of the individuals had an influence on final CT_max_ and on the change in CT_max_ (p-value = 0.03 in both cases, data not shown).

| Coefficients | Estimate | Std. Error | t-value | p-value |
| --- | --- | --- | --- | --- |
| *(a) glm(Basal CT_max_ ~ Group + Sex + SVL + Mass, family = “gaussian”)* ***AIC: 117.14*** | | | | |
| (Intercept) | 65.675 | 7.296 | 9.001 | **7.9 x 10^-9^** |
| Group (3h) | 0.879 | 0.797 | 1.102 | 0.282 |
| Group (6h) | -0.427 | 0.801 | -0.534 | 0.598 |
| Sex (M) | 1.145 | 0.717 | 1.596 | 0.124 |
| SVL (cm) | -2.529 | 0.776 | -3.259 | **0.00359** |
| Mass (g) | 0.093 | 0.068 | 1.369 | 0.184 |
| Null deviance: 107.057 on 27 df; Residual deviance: 65.221 on 22 df | | | | |
| *(b) glm(Final CT_max_ ~ Group*Basal CT_max_ + Sex + SVL + Mass, family = “gaussian”)*  ***AIC: 94.176*** | | | | |
| (Intercept) | 11.575 | 12.051 | 0.961 | 0.348 |
| Group (3h) | 47.594 | 11.035 | 4.313 | **0.000375** |
| Group (6h) | 13.459 | 11.836 | 1.137 | 0.269 |
| Basal CT_max_ | 0.855 | 0.201 | 4.249 | **0.000434** |
| Sex (M) | 0.563 | 0.489 | 1.153 | 0.263 |
| SVL (cm) | -0.120 | 0.613 | -0.197 | 0.846 |
| Mass (g) | -0.097 | 0.047 | -2.043 | 0.055 |
| Group(3h):Basal CT_max_ | -1.131 | 0.266 | -4.243 | **0.000440** |
| Group(6h):Basal CT_max_ | -0.313 | 0.291 | -1.079 | 0.294 |
| Null deviance: 83.170 on 27 df, Residual deviance: 23.184 on 19 df | | | | |
| *(c) glm(Change in CT_max_ ~ Group*Basal CT_max_ + Sex + SVL + Mass, family = “gaussian”)* ***AIC = 94.176*** | | | | |
| (Intercept) | 11.575 | 12.051 | 0.961 | 0.348 |
| Group (3h) | -7.162 | 11.035 | -0.649 | 0.524 |
| Group (6h) | 13.459 | 11.836 | 1.137 | 0.269 |
| Basal CTmax | -0.144 | 0.201 | -0.718 | 0.481 |
| Sex (M) | 0.563 | 0.489 | 1.153 | 0.263 |
| SVL (cm) | -0.120 | 0.613 | -0.197 | 0.846 |
| Mass (g) | -0.097 | 0.047 | -2.043 | 0.055 |
| Group(3h):Basal CT_max_ | 0.182 | 0.266 | 0.684 | 0.502 |
| Group(6h):Basal CT_max_ | -0.313 | 0.291 | -1.079 | 0.294 |
| Null deviance: 40.735 on 27 df, Residual deviance: 23.184 on 19 df | | | | |

**2. Testing of Assumptions for Statistical Analyses**

**2.1 Heating Rates**

The differences between basal heating rates and final heating rates across all time-interval treatment groups for all individuals were found to be normally distributed using a Shapiro-Wilks test (W = 0.9799, p-value = 0.848) and have equal variance across all time-interval treatment groups using a Levene’s test (df = 2, F-value = 1.127, p-value = 0.3399). Differences between basal heating rates and final heating rates of all individuals were also found to be normally distributed within each time-interval treatment group (3h: W = 0.95268, p-value = 0.7195; 6h: W = 0.97593, p-value = 0.9401; 24h: W = 0.95805, p-value = 0.7634).

**2.2 Basal CT_max_, final CT_max_, and change in CT_max_**

Basal CT_max_ (W=0.93661, p-value = 0.09062), final CT_max_ (W=0.95448, p-value = 0.2562), and change in CT_max_ (W = 0.97691, p-value = 0.771) of all individuals for all treatment groups were found to be normally distributed after performing a Shapiro-Wilks tests using the function “shapiro.test” function from the *stats* package. To test for homogeneity of variance across time-interval treatment groups, Levene’s tests were performed on basal CT_max_, final CT_max_, and change in CT_max_ for all individuals for all time-interval treatment groups using the “leveneTest” function from the *car* package. Variance across time-interval treatment groups was found to be equal for basal CT_max_ (df = 2, F-value = 0.1092, p-value = 0.897), final CT_max_ (df = 2, F-value = 0.2488, p-value = 0.7817), and change in CT_max_ (df = 2, F-value = 0.649, p-value = 0.5311). From each of the GLM models which included the mass of all individuals measured before basal CT_max_ experiments, the residuals for basal CT_max_ (W = 0.97046, p-value = 0.593), final CT_max_ (W = 0.93022, p-value = 0.06243), and change in CT_max_ (W = 0.93022, p-value = 0.06243) were all found to be normally distributed based on a Shapiro-Wilks test. The residuals for the same GLM models for basal CT_max_ (W = 0.96357, p-value = 0.4222) and final CT_max_ (W = 0.98193, p-value = 0.8938) with only significant variables included in each model were normally distributed. The GLM models for final CT_max_ (W = 0.9375, p-value = 0.1057) and change in CT_max_ (W = 0.9375, p-value = 0.1057) which included the mass of all individuals measured before final CT_max_ experiments were also found to be normally distributed from the Shapiro-Wilks test.Residuals for the same GLM models for final CT_max_ (W = 0.94903, p-value = 0.2031) and change in CT_max_ (W = 0.96079, p-value = 0.3852) including only significant variables for each model were found to be normally distributed. Levene’s tests also found equal variances across all time-interval treatment groups for all residuals for basal CT_max_ (df = 2, F-value = 0.2786, p-value = 0.7592), final CT_max_ (df = 2, F-value = 1.8554, p-value = 0.1773), and change in CT_max_ (df = 2, F-value = 1.8554, p-value = 0.1773) for the GLM models which included the mass of all individuals measured before basal CT_max_ experiments. The residuals for the same GLM models for basal CTmax (df = 2, F-value = 0.6499, p-value = 0.5307) and final CT_max_ (df = 2, F-value = 0.924, p-value = 0.41) with only significant variables included each model were found to be normally distributed. Equal variances were found across all time-interval groups for all residuals for final CT_max_ (df = 2, F-value = 2.0425, p-value = 0.1517) and change in CT_max_ (df = 2, F-value = 2.0425, p-value = 0.1517) for GLM models which included the mass of all individuals measured before final CT_max_ experiments. Residuals for the same GLM models for final CT_max_ (df = 2, F-value = 2.672, p-value = 0.0896) and change in CT_max_ (df = 2, F-value = 1.6015, P-value = 0.2224) only including significant variables in each model were found to be normally distributed.
